# Loss of histone methyltransferase SETD1B in oogenesis results in the redistribution of genomic histone 3 lysine 4 trimethylation

**DOI:** 10.1101/2021.03.11.434836

**Authors:** Courtney W. Hanna, Jiahao Huang, Christian Belton, Susanne Reinhardt, Andreas Dahl, Simon Andrews, A. Francis Stewart, Andrea Kranz, Gavin Kelsey

## Abstract

Histone 3 lysine 4 trimethylation (H3K4me3) is an epigenetic mark found at gene promoters and CpG islands. H3K4me3 is essential for mammalian development, yet mechanisms underlying its genomic targeting are poorly understood. H3K4me3 methyltransferases SETD1B and MLL2 are essential for oogenesis. We investigated changes in H3K4me3 in *Setd1b* conditional knockout (cKO) oocytes using ultra-low input ChIP-seq, with comparisons to DNA methylation and gene expression analyses. H3K4me3 was redistributed in *Setd1b* cKO oocytes showing losses at active gene promoters associated with downregulated gene expression. Remarkably, many regions also gained H3K4me3, in particular those that were DNA hypomethylated, transcriptionally inactive and CpG-rich, which are hallmarks of MLL2 targets. Consequently, loss of SETD1B disrupts the balance between MLL2 and *de novo* DNA methyltransferases in determining the epigenetic landscape during oogenesis. Our work reveals two distinct, complementary mechanisms of genomic targeting of H3K4me3 in oogenesis, with SETD1B linked to gene expression and MLL2 to CpG content.

**Graphical Abstract:** In oogenesis, SETD1B and CXXC1 target H3K4me3 to actively transcribed gene promoters, while MLL2 targets transcriptionally inactive regions based on underlying CpG composition (upper panel). When SETD1B is ablated, H3K4me3 is lost at a subset of active promoters, resulting in downregulation of transcription (lower panel). Loss of SETD1B alters the activity of MLL2, permitting MLL2 to deposit H3K4me3 at CpG-rich regions, many of which should otherwise be DNA methylated. Thus, it is evident that MLL2 and de novo DNMTs compete for genomic occupancy late in oogenesis, and loss of SETD1B disrupts the balance of these mechanisms.

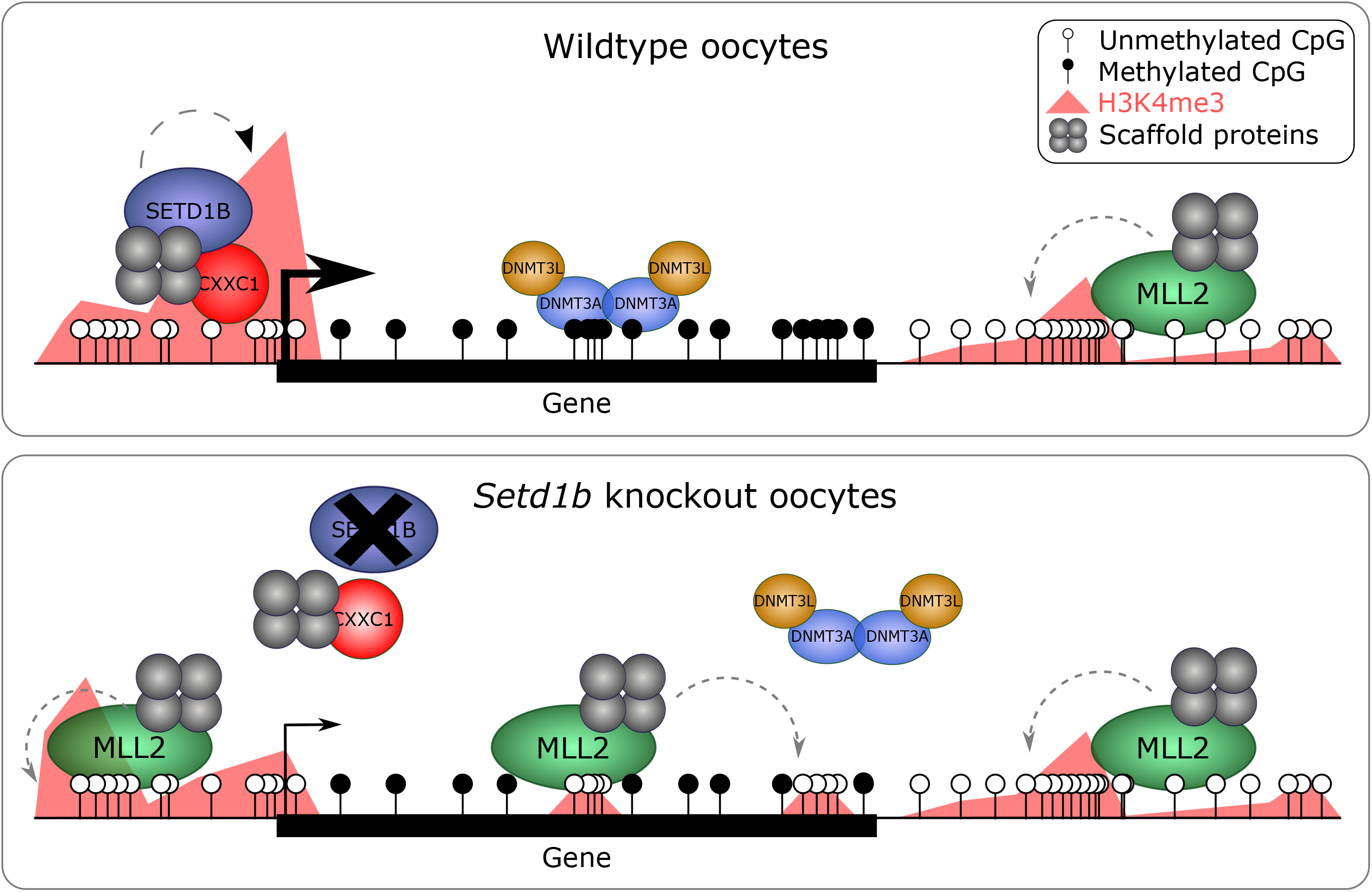

## Introduction

Histone 3 lysine 4 trimethylation (H3K4me3) is a hallmark of active gene promoters and is amongst the most conserved eukaryotic epigenetic modifications (1, 2, 3). Despite high concordance with promoter activity, the function of H3K4me3 in gene regulation remains enigmatic (4). Loss of H3K4me3 has surprisingly little impact on global transcription in several contexts (5, 6). In mammalian cells, H3K4me3 is not only correlated with gene transcription, but is strongly associated with CpG islands irrespective of transcriptional activity (7). Some studies indicate a role for H3K4me3 in transcriptional induction (8) as well as the maintenance of transcriptional activity (9). However, H3K4me3 has been paradoxically associated with gene repression in yeast (5, 10) and global transcriptional silencing in mouse oocytes (11). In contrast to doubts regarding its role in the regulation of transcription, H3K4me3 serves to prevent the imposition of repression either by Polycomb-Group action (12) or by DNA methylation, as it inhibits binding of *de novo* DNA methyltransferases (DNMTs) at gene promoters and CpG islands (13, 14). Indeed, before zygotic genome activation in zebrafish development, H3K4 methylation appears to prevent DNA methylation rather than serving a direct role in transcription (15).

Due to the importance of H3K4me3 in development, fertility and disease, there is a need to understand the mechanisms that govern the placement and maintenance of H3K4me3 in chromatin. Several mechanisms that do not rely on positioning by sequence-specific DNA binding proteins (i.e., transcription factors) have been identified. In yeast, the sole H3K4 methyltransferase, Set1 (10, 16), associates with both serine 5 and serine 2 phosphorylated forms of elongating RNA polymerase II (17) and the size of the H3K4me3 promoter peak closely correlates with mRNA production indicating that H3K4me3 deposition is a consequence of transcription (18, 19). The yeast Set1 complex, termed either Set1C or COMPASS, is highly conserved in evolution (16, 20, 21, 22). It includes a subunit involved in chromatin targeting, Spp1 (in mammals, CXXC1 or CFP1), which contains a PHD finger that binds to H3K4me3 (8, 14, 23, 24), and potentially directs Set1C binding to active promoters. CXXC1, which is a subunit of the mammalian SETD1A and SETD1B complexes, also includes the CxxC zinc finger that binds the other hallmark of mammalian promoters - unmethylated CpG dinucleotides (7, 25, 26).

In mammals, the mechanisms that target H3K4me3 in chromatin have been additionally challenging to disentangle because mammals have six partially redundant H3K4 methyltransferase enzymes (SETD1A, SETD1B, MLL1, MLL2, MLL3, and MLL4) (22, 27, 28, 29). Each of these H3K4 methyltransferases forms a complex with a common set of four scaffold proteins: WDR5, RbBP5, ASH2L, and DPY30 (16, 22, 30, 31, 32). Structures of these highly conserved quintets have recently been solved (33). Although CXXC1 is only found in the SETD1A and SETD1B complexes, MLL1 and MLL2 include the same combination of a PHD finger that binds H3K4me3 and a CxxC zinc finger that binds unmethylated CpG dinucleotides. Hence, at least four of the six complexes can potentially bind to CpG island promoters without reliance on guidance by a DNA sequence-specific transcription factor. Despite this insight, we still have a poor understanding of the differential recruitment of the six H3K4 methyltransferases across genomic regions and in differing cell types. Furthermore, we are far from understanding the distinct functional specificities of the six mammalian H3K4 methyltransferases (27, 29, 34, 35, 36).

Murine oogenesis presents a unique window into the epigenetic mechanisms that reset the genome for launching the developmental program, and this resetting critically relies on H3K4 and DNA methylation. In the oocyte, our previous work established that SETD1A, MLL1 and MLL3 are not required and only SETD1B, MLL2 and at least one more as yet unidentified H3K4 methyltransferase are required (11, 27, 37). CXXC1 is essential for oogenesis, presumably to support SETD1B function, given the similarities in phenotypes of oocyte-specific *Cxxc1* and *Setd1b* knockouts (37, 38, 39). We recently demonstrated the power of ultra-low input native ChIP-seq (ULI-nChIP-seq) in *Mll2* conditional knockout (cKO) oocytes to elucidate the changes in the H3K4me3 landscape (40). Specifically, we identified that MLL2 was responsible for deposition of H3K4me3 at transcriptionally silent, unmethylated genomic regions, dependent on the underlying CpG density, thus giving rise to the characteristic broad domains of H3K4me3 in mouse oocytes (41, 42), but had relatively little impact on the oocyte transcriptome. Conditional deletion of *Setd1b* in oocytes altered the oocyte transcriptome without changing H3K4me3 abundance as evaluated by immunofluorescence (37). Loss of SETD1B in oogenesis results in increased atretic follicles, meiotic spindle defects and zona pellucida defects, and upon fertilisation, failure to progress past the 2-cell stage (37). Having established that MLL2 deposits the majority of H3K4me3 across the oocyte genome shortly before global transcriptional silencing, here we address the role of SETD1B in establishing the oocyte H3K4me3 landscape using ULI-nChIP-seq.

## Materials and Methods

### Sample collection

Experiments were performed in accordance with German animal welfare legislation, and were approved by the relevant authorities, the Landesdirektion Dresden.

We conditionally mutated *Setd1b* during oocyte development by crossing the *Setd1b* conditional line with the *Gdf9*-Cre line, which excises from early folliculogenesis onwards (43). The *Setd1b* conditional line as well as the Cre deleter are maintained in C57BL/6OlaHsd background. To obtain *Setd1b*^*FD*/+;^*Gdf9*-Cre males, *Setd1b*^*FD*/FD^ females were crossed with Gdf9-Cre males. Then these males were crossed to *Setd1b*^*FD*/FD^ females resulting in offspring with the genotype *Setd1b*^*FDC*/FDC^;*Gdf9*-Cre (*Setd1b* cKO females) and *Setd1b*^*FD*/FD^ or *Setd1b*^*FDC*/+^;*Gdf9*-Cre (*Setd1b* WT females) (37).

Ovaries were collected from 21-day-old females and digested with 2mg/mL collagenase (Sigma) and 0.02% trypsin solution, agitated at 37°C for 30 min. GV oocytes were collected in KSOM medium (Millipore) and washed with PBS. Four GV oocytes for each genotype were collected for single cell RNA-seq (scRNA-seq). Approximately 100 GV oocytes were collected for each of three biological replicates of each genotype for post-bisulphite adaptor tagging (PBAT). Approximately 200 GV oocytes were collected for each of four biological replicates of each genotype for H3K4me3 ULI-nChIP-seq. All molecular experiments were performed as previously described (40), but are described in brief below.

### Preparation of scRNA-seq libraries

Cells were lysed and RNA was reverse transcribed and amplified according to the SMARTer Ultra Low RNA Kit for Illumina Sequencing (version 1, Clontech). Libraries were prepared from cDNA using the NEBNext Ultra DNA library preparation for Illumina sequencing with indexed adaptors (New England Biolabs). Library preparation failed for one replicate of *Setd1b* WT. Libraries were multiplexed for 75-bp single-read sequencing on an Illumina HiSeq 2000.

### Preparation of PBAT libraries

Cells were lysed with 0.5% SDS in EB buffer and bisulphite treated with the Imprint DNA Modification kit (Sigma). The resulting DNA was purified using columns and reagents from the EZ DNA Methylation Direct kit (Zymo Research). First-strand synthesis was performed with Klenow Exo-enzyme (New England Biolabs) using a customized biotin-conjugated adaptor containing standard Illumina adaptor sequences and 9 bp of random sequences (9N), as previously described (40). Following exonuclease I (New England Biolabs) treatment and binding to Dynabeads M-280 Streptavidin beads (Thermo Fisher Scientific), second-strand synthesis was performed with Klenow Exo-enzyme (New England Biolabs) using a customized adaptor containing standard Illumina adaptor sequences and 9 bp of random sequences. Ten PCR cycles with Phusion High-Fidelity DNA polymerase (New England Biolabs) were used for library amplification with indexed adaptors. Libraries were multiplexed for 100-bp paired-end sequencing on an Illumina HiSeq 2500.

### Preparation of ULI-nChIP-seq libraries

The ULI-nChIP-seq protocol was performed using our previous protocol (40), which incorporates several adaptions from the original published protocol (44). Samples were thawed on ice and permeabilized using nuclei EZ lysis buffer (Sigma) and 0.1% Triton-X-100/0.1% deoxycholate. Micrococcal nuclease digestion was completed with 200 U of micrococcal nuclease (New England Biolabs) at 21°C for 7.5 min. Chromatin samples were precleared in complete immunoprecipitation buffer with Protein A/G beads rotating for 2 hours at 4°C. For each sample, 125ng of anti-H3K4me3 antibody (Diagenode, C15410003) was bound to Protein A/G beads in complete immunoprecipitation buffer for 3 hours rotating at 4°C. Chromatin was then added to the antibody-bound beads and samples were rotated overnight at 4°C. Chromatin-bound beads were washed with two low-salt washes and one high-salt wash and DNA was then eluted from the beads at 65°C for 1.5 hours. Eluted DNA was purified with solid-phase reversible immobilization (SPRI) purification with Sera-Mag carboxylate-modified Magnetic SpeedBeads (Fisher Scientific) at a 1.8:1 ratio. Library preparation was completed using the MicroPlex Library Preparation kit v2 (Diagenode) with indexed adaptors, as per the manufacturers guidelines. Libraries were multiplexed for 75 bp Single End sequencing on an Illumina NextSeq500.

### Library mapping and trimming

Fastq sequence files were quality and adaptor trimmed with trim galore v0.4.2 using default parameters. For PBAT libraries, the -clip option was used. Mapping of ChIP-seq data was performed with Bowtie v2.2.9 against the mouse GRCm38 genome assembly. The resulting hits were filtered to remove mappings with a MAPQ scores < 20. Mapping of RNA-seq data was performed with Hisat v2.0.5 against the mouse GRCm38 genome assembly, as guided by known splice sites taken from Ensembl v68. Hits were again filtered to remove mappings with MAPQ scores <20. Sequencing depths for all libraries are provided in Supplementary Table 1.

Mapping and methylation calling of bisulphite-seq data was performed using Bismark v0.16.3 in PBAT mode against the mouse GRCm38 genome assembly. Trimmed reads were first aligned to the genome in paired-end mode to be able to detect and discard overlapping parts of the reads while writing out unmapped singleton reads; in a second step remaining singleton reads were aligned in single-end mode. Alignments were carried out with Bismark (45) with the following set of parameters: a) paired-end mode: --pbat; b) single-end mode for Read 1: --pbat; c) single-end mode for Read 2: defaults. Reads were then deduplicated with deduplicate_bismark selecting a random alignment for positions that were covered more than once. Following methylation extraction, CpG context files from PE and SE runs were used to generate a single coverage file (the “Dirty Harry” procedure).

### Publicly available datasets

Publicly available datasets were downloaded from Gene Expression Omnibus (GEO), including: H3K27me3 and H3K27ac ChIP-seq in GV oocytes, H3K4me3 ChIP-seq from d5 non-growing oocytes, d10 growing oocytes, d15 GV oocytes, d25 GV oocytes, *Dnmt3* cDKO and *Dnmt3* WT GV oocytes, and RNA-seq from *Mll2* cKO and WT GV oocytes (GSE93941) (40), RNA-seq for GV oocytes (GSE70116) (46), RNA-seq from size-selected oocytes (GSE86297) (47), RNA-seq from *Setd1b* cKO and WT MII oocytes (GSE85360) (37), RNA-seq from *Cxxc1* cKO and WT GV oocytes (GSE85019) (39), H3K4me3 ChIP-seq and bisulphite-seq from *Cxxc1* cKO and WT GV oocytes (GSE159581) (48), and bisulphite-seq from d12 and d15 GV oocytes (GSE72784) (41, 42). All raw data files were mapped and trimmed using the pipelines described above.

### ChIP-seq analysis

5kb running windows were used for quantitative analysis of H3K4me3 ChIP-seq data using RPKM, with mapping artefacts excluded from analysis (RPKM>4 in any input sample). Poor quality H3K4me3 ChIP-seq libraries were excluded from analysis, which included one replicate of *Setd1b* WT and one replicate of *Setd1b* cKO. Poor enrichment was defined as a cumulative distribution plot reflecting a signal-to-noise similar to input and presenting as an outlier in hierarchical clustering. For all subsequent analyses, including the use of publically available H3K4me3 ChIP-seq data, valid 5kb windows were quantitated using enrichment normalisation of RPKM in SeqMonk, as previously described (40). LIMMA statistic (p<0.05 corrected for multiple comparisons) was used to identify differentially enriched H3K4me3 peaks between *Setd1b* cKO and WT GV oocytes, using an average normalised RPKM for replicates within each group. Overlapping TSSs and promoters (TSS +/-500bp) were classified as previously described (40). In brief, active promoters were defined as those marked with H3K27ac or transcripts with >1 FPKM in GV oocytes, weak promoters were defined those transcripts with an FPKM between 0.1 and 1, and inactive promoters were defined as those transcripts with undetectable gene expression (FPKM<0.1), fell within a DNA methylated domain in GV oocytes (46) or marked with H3K27me3.

### RNA-seq analysis

RNA-seq datasets were quantitated using the RNA-seq quantitation pipeline in Seqmonk, over previously defined oocyte transcripts (46). Transcript isoforms were merged for all analyses. A correlation matrix between replicates is provided in Supplementary Table 2. Differentially expressed genes (DEGs) were identified in three comparisons using DESeq2 (p<0.05 corrected for multiple comparisons) and a >2-fold change in expression: (1) *Setd1b* cKO and WT GV oocytes, (2) 10-30μm oocytes (non-growing oocytes) and GV oocytes, and (3) *Cxxc1* cKO and WT GV oocytes. Fold enrichment for genes upregulated in oogenesis and DEGs among the *Setd1b, Cxxc1* and *Mll2* cKOs was calculated relative to all oocyte transcripts and increase in observed over expected overlap between DEGs from different conditions was statistically compared using Chi-square. Gene ontology analysis was done using DAVID (https://david.ncifcrf.gov/) for up- and down-regulated DEGs, using the default settings with the addition of the UP_TISSUE category.

### Sequence composition analysis

Dimer composition of candidate 5kb windows (gains and losses of H3K4me3 in *Setd1b* cKO and a random set) was assessed using compter (https://www.bioinformatics.babraham.ac.uk/projects/compter/). The mouse genome was used as the background and values were expressed as log2 observed/expected. Enrichment values for each dimer in each condition were summarised by taking the mean, and the difference in mean log ratio (*Setd1b* cKO - Random) was calculated and plotted.

### DNA methylation analysis

DNA methylation datasets were analysed using 100-CpG running windows, with a minimum coverage of at least 10-CpGs, using the bisulphite-seq quantitation pipeline in SeqMonk. Differentially methylated regions (DMRs) were identified between *Setd1b* cKO and WT GV oocytes using logistic regression (p<0.05 corrected for multiple comparisons) and a >20% difference in methylation.

## Results

### Loss of SETD1B in oogenesis results in gains and losses of H3K4me3

In order to evaluate the role of SETD1B in the H3K4me3 landscape in oocytes, we isolated fully-grown germinal vesicle (GV) oocytes from the ovaries of 21-day old mice after oocyte-specific ablation of *Setd1b* driven by *Gdf9*-Cre (37, 43). Pools of approximately 200 oocytes were collected and processed for ULI-nChIP-seq (40). The genome-wide distribution of H3K4me3, assessed using 5kb running windows, differed only modestly between *Setd1b* cKO (N=3) and *Setd1b* WT (N=3) GV oocytes (Figure 1A, 1B, Supplementary Figure 1A, 1B), in concordance with the observations made by immunofluorescence (37). However, the biological replicates for *Setd1b* cKO separated from WT in a hierarchical cluster (Supplementary Figure 1A), and we identified 3.6% of the genome that exhibited significant gains or losses of H3K4me3 (Figure 1A, 1B). The majority of differentially enriched windows were not at gene promoters (Supplementary Figure 1C). However, given the potential role of H3K4me3 in gene regulation, we focused first on gene promoters, considering active, weak and inactive promoters, as previously described by histone modifications and gene expression in GV oocytes (40). Promoter-associated 5kb windows that lost H3K4me3 in *Setd1b* cKO oocytes were significantly enriched at active gene promoters, defined by either H3K27ac or high gene expression (Figure 1C). Conversely, promoter-associated 5kb windows that gained H3K4me3 in the *Setd1b* cKO oocytes were significantly enriched at inactive promoters, defined by either the presence of H3K27me3, DNA methylation or undetectable gene expression (Figure 1C). This trend was recapitulated when we looked genome-wide at enrichment for H3K27ac and H3K27me3 (Supplementary Figure 1C, 1D).

**Figure 1.**
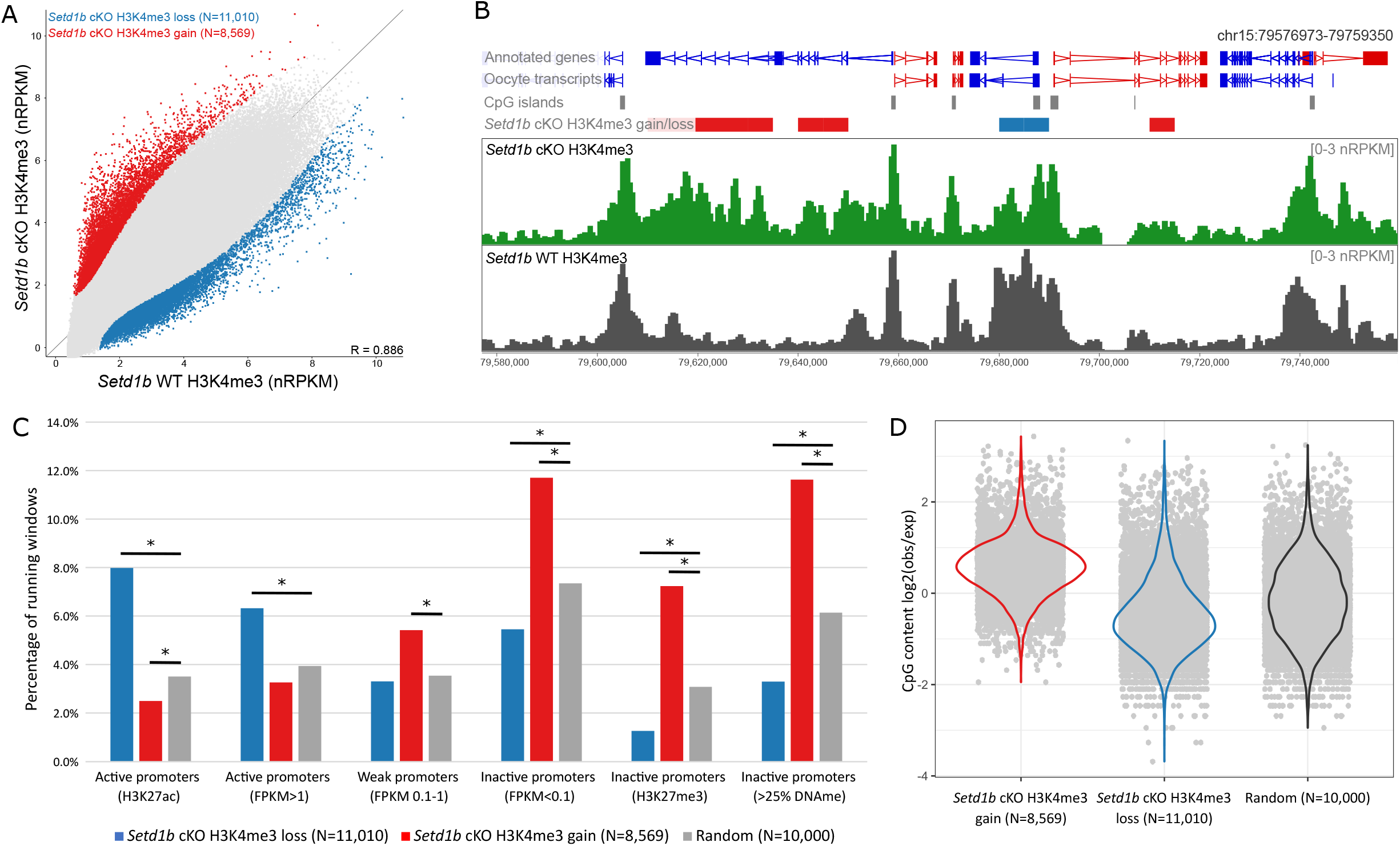
**(A)** The scatterplot shows average normalised enrichment for H3K4me3 for 5kb running windows (N=544,879) between *Setd1b* cKO (N=3) and WT (N=3) d21 GV oocytes. Differentially enriched windows were identified using LIMMA statistic (p<0.05, corrected for multiple comparisons), and those that show a loss in *Setd1b* cKO are shown in blue and a gain in red. **(B)** The screenshot shows the normalised enrichment for H3K4me3 for 1kb running windows with a 500bp step in *Setd1b* cKO and WT GV oocytes. Significant differentially enriched 5kb windows are shown in the *Setd1b* gain/loss track, with red and blue bars showing windows that gain and lose H3K4me3 in *Setd1b* cKO, respectively. **(C)** The barplot shows the overlap between sets of promoters and differentially enriched windows that gain or lose H3K4me3 in the *Setd1b* cKO compared to a random set of 5kb running windows. Pairwise Chi-Square statistic was used to compare each set of enriched windows to random. The asterisk shows those comparisons that were significant after Bonferroni correction for multiple comparisons (p<0.008). **(D)** The beanplot shows the enrichment for CG content among the 5kb running windows that gain or lose H3K4me3 in *Setd1b* cKO oocytes compared to a random set of 5kb windows.

MLL2 is responsible for the majority of transcription-independent H3K4me3 in the oocyte (40), so this finding suggests that there may be increased MLL2 action upon loss of SETD1B. Because MLL2 is highly dependent on underlying CpG content (40), we then examined sequence composition in the regions that gain H3K4me3 in the *Setd1b* cKO and found a highly significant enrichment for CpG content compared to a random set of regions (Figure 1D, Supplementary Figure 1E). Furthermore, we observed a significant depletion of CpG content at regions that lose H3K4me3 in *Setd1b* cKO (Figure 1D). Together these findings indicate that the loss of SETD1B in oocytes results in a redistribution of H3K4me3, away from active and CpG-poor regions to repressed and CpG-rich regions of the genome.

### Loss of promoter H3K4me3 is linked to downregulated gene expression of oocyte transcriptional regulators in Setd1b cKO oocytes

We investigated whether the changes in promoter H3K4me3 in the *Setd1b* cKO oocytes were associated with changes in gene expression. Using single-cell RNA-seq, we identified 1519 differentially expressed genes (DEGs) between *Setd1b* cKO (N=4) and *Setd1b* WT (N=3) GV oocytes, with 594 genes downregulated and 925 gene upregulated in the *Setd1b* cKO (Figure 2A, Supplementary Figure 2A). We then assessed DEG promoters and found that H3K4me3 was significantly lower in *Setd1b* cKO compared to *Setd1b* WT GV oocytes at promoters of downregulated DEGs (p<0.0001, two-tailed t-test), while there was no difference at upregulated DEGs (p=1.0, twotailed t-test) (Figure 2B). Nevertheless, oocyte promoters that showed a significant gain or loss of H3K4me3 also exhibited an up- or down-regulation of gene expression, respectively (Supplementary Figure 2B), reinforcing the relevance of the association.

**Figure 2.**
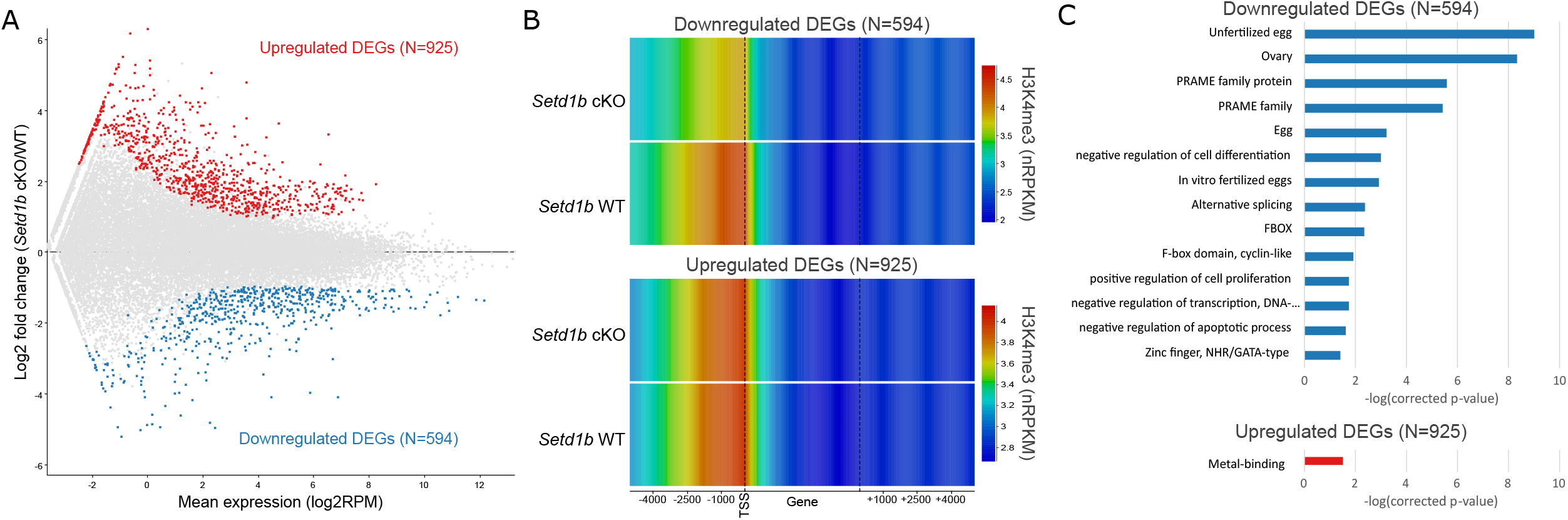
**(A)** The MA plot shows the expression of oocyte transcripts (N=33,437) in *Setd1b* cKO and WT GV oocytes as measured by single-cell RNA-seq. Differentially expressed genes (DEGs) were identified using DESeq2 (p<0.05 corrected for multiple comparisons) and a >2-fold change in expression. **(B)** The heatmap shows H3K4me3 enrichment (normalised RPKM) in *Setd1b* cKO and WT oocytes across downregulated (upper panel) and upregulated (lower panel) DEGs, including 5kb upstream and downstream. **(C)** The barplot shows the –log(corrected p-value) for significant gene ontology categories for down- and up-regulated DEGs in *Setd1b* cKO GV oocytes.

Gene ontology of down- and up-regulated DEGs showed a significant enrichment for the egg, ovary and transcriptional regulators among downregulated DEGs (Figure 2C). Compared to published RNA-seq data from *Setd1b* cKO MII oocytes (37), *Setd1b* cKO DEGs showed consistent changes in expression, despite the distinctive transcriptomes between GV and MII oocytes (Supplementary Figure 2C, 2D). Importantly, no difference in expression of any of the other H3K4 methyltransferases in *Setd1b* cKO GV oocytes was apparent (Supplementary Figure 2E). Together, these data suggest that downregulated expression of oocyte transcriptional regulators in *Setd1b* cKO oocytes may be a consequence of loss of SETD1B-dependent promoter H3K4me3.

### SETD1B/CXXC1-deposited H3K4me3 is associated with transcriptional upregulation of a subset of oogenesis genes

Active promoters display H3K4me3 in primary non-growing oocytes (40). Because the *Gdf9*-Cre recombinase used to generate *Setd1b* cKO oocytes is active from postnatal day 3 (43), it is likely that only *de novo* and renewed H3K4me3 deposited by SETD1B from this stage onward will be affected and the established, early stage, H3K4me3 will persist. Therefore, we further focused on gene expression and H3K4me3 changes at transcripts that are upregulated during oogenesis (47). There was a significant enrichment of genes upregulated during oogenesis among downregulated DEGs (p<0.0001, Chi-square) but not upregulated DEGs (p=0.1, Chi-square) in *Setd1b* cKO oocytes (Figure 3A, Supplementary Figure 3A).

**Figure 3.**
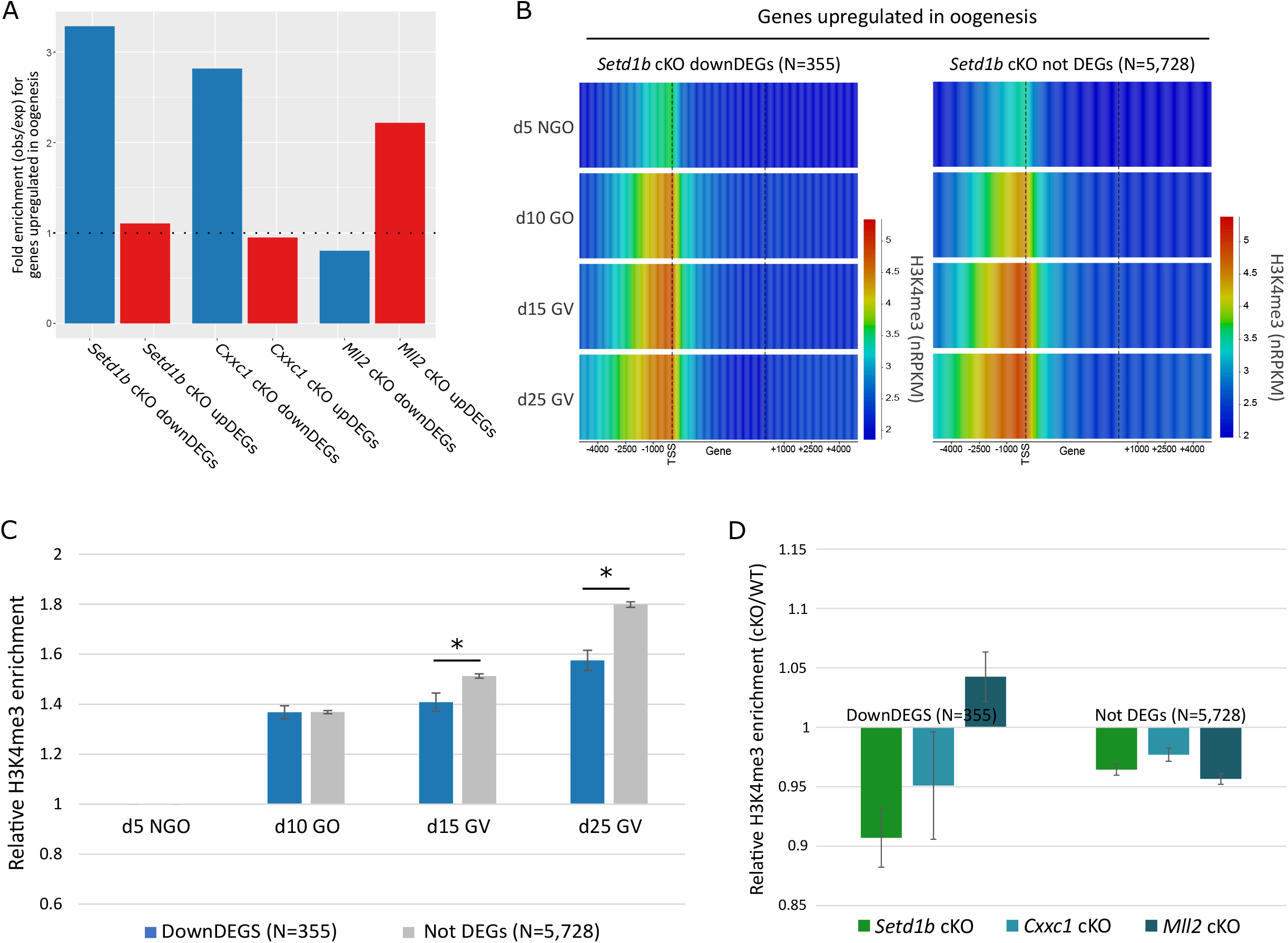
**(A)** The barplot shows the fold enrichment for genes that are upregulated in oogenesis (N=6,083) among the *Setd1b* cKO down- and upregulated DEGs (p<0.0001 and p=0.1, respectively), *Cxxc1* cKO down- and upregulated DEGs (p<0.0001 and p=0.2, respectively) and *Mll2* cKO down- and upregulated DEGs (p=0.05 and p<0.0001, respectively), relative to all oocyte transcripts. **(B)** The heatmap shows H3K4me3 enrichment (normalised RPKM) in d5 non-growing oocytes (NGOs), d10 growing oocytes (GOs), d15 GV, d25 GV oocytes across transcripts that are upregulated during oogenesis, including 5kb upstream and downstream of the gene. The left panel shows genes that are significantly downregulated in *Setd1b* cKO (downDEGs), while the right panel shows genes that show no significant expression change in *Setd1b* cKO (not DEGs). Promoter H3K4me3 enrichment was statistically compared using one-way ANOVA (p<0.0001) for the overlapping 5kb windows. **(C)** The barplot shows the average relative increase in H3K4me3 enrichment for 5kb windows overlapping promoters of downDEGs and not DEGs in d10 GOs, d15 GV and d25 GV oocytes compared to d5 NGOs. Pair-wise comparisons were done using a two-tailed t-test. The asterisk shows significant p-values (p<0.01). Error bars show standard error of the mean. **(D)** The barplot shows the relative H3K4me3 enrichment for *Setd1b* cKO, *Cxxc1* cKO and *Mll2* cKO over matched WT controls for 5kb windows overlapping promoters of downDEGs and not DEGS. Relative H3K4me3 enrichment was statistically compared using one-way ANOVA for downDEGs (p<0.01) and not DEGs (p=0.027). Error bars show standard error of the mean.

As CXXC1 and SETD1B function together in the SETD1B complex (49) and have similar phenotypes when ablated in oocytes (37, 38, 39), we compared gene expression changes in *Setd1b* cKO with publicly available data for *Cxxc1* cKO GV oocytes (39). Similar to the *Setd1b* cKO DEGs, we observed a significant enrichment for genes upregulated during oogenesis among downregulated DEGs in the *Cxxc1* cKOs (p<0.0001, Chi-square), but not upregulated DEGs (p=0.2, Chi-square) (Figure 3A, Supplementary Figure 3B). We see a more than 10-fold enrichment for *Cxxc1* cKO downregulated DEGs among downregulated DEGs identified in the *Setd1b* cKO oocytes (p<0.0001) (Supplementary Figure 3C). There is a significant correlation between fold change in expression in *Setd1b* cKO and *Cxxc1* cKO for *Setd1b* DEGs (Supplementary Figure 3D). These data indicate that CXXC1 facilitates SETD1B targeting to actively transcribed promoters, and that the corresponding SETD1B-deposited H3K4me3 contributes to, or is a consequence of, upregulation of gene expression during oogenesis for at least a subset of genes.

To further evaluate why only a subset of upregulated genes was impacted in *Setd1b* cKO oocytes, we evaluated the dynamics of promoter H3K4me3 for genes upregulated in oogenesis that were downregulated DEGs compared to genes not called as differentially expressed (not DEGs) in *Setd1b* cKO oocytes (N=355 and 5728, respectively). Promoters of *Setd1b* cKO downregulated DEGs gained the majority of H3K4me3 early in oogenesis (between d5 and d10), while the promoters of genes that were not DEGs continued to gain H3K4me3 throughout oogenesis until d25 (Figure 3B, 3C). Therefore, we hypothesized that H3K4me3 these sets of promoters may have differing dependencies on H3K4 methyltransferases. To test this, we included publically available H3K4me3 ChIP-seq data for *Cxxc1* cKO and *Mll2* cKO GV oocytes (40, 48). Globally, *Setd1b* cKO and *Cxxc1* cKO GV oocytes show remarkably similar patterns of H3K4me3 (Supplementary Figure 4A, 4B, 4C), further supporting their co-dependence in the oocyte. Notably, promoters of *Setd1b* cKO downregulated DEGs lost H3K4me3 in *Setd1b* and *Cxxc1* cKO, but not *Mll2* cKO oocytes, whereas promoters of not DEGs showed reduced H3K4me3 in all three cKOs (Figure 3D). This finding indicates that the majority of genes upregulated in oogenesis show a partial redundancy between SETD1B/CXXC1 and MLL2 for H3K4me3 deposition at their promoters. However, at a subset of promoters, H3K4me3 appears to be solely reliant on SETD1B/CXXC1 and ablation of either leads to a failure to appropriately induce gene expression during oogenesis.

### Setd1b KO oocytes show widespread gains of H3K4me3 across regions that lose DNA methylation

H3K4me3 and DNA methylation are mutually exclusive in the oocyte, despite both having atypical genomic patterning compared to somatic cells. In the oocyte, DNA methylation is almost exclusively restricted to transcribed gene bodies (50, 51), whereas H3K4me3 forms broad domains across much of the remaining unmethylated fraction of the genome (41, 42) through the activity of MLL2 (40). To evaluate the relationship between DNA methylation and the regions that show altered H3K4me3 in *Setd1b* cKO oocytes (N=8,569), we assessed genome-wide patterns of DNA methylation using post-bisulphite adaptor tagging (PBAT). DNA methylation in *Setd1b* cKO and WT GV oocytes was distinct (Figure 4A), with the vast majority (94%) of differentially methylated regions (DMRs) losing DNA methylation in the *Setd1b* cKOs (Figure 4B). This finding is not explained by a difference in mRNA expression of the *de novo Dnmts*, which are unchanged in *Setd1b* cKO GV oocytes (Supplementary Figure 2E). A similar loss of DNA methylation was also observed in *Cxxc1* cKO oocytes (Supplementary Figure 5A, 5B, 5C). Strikingly, we observed a significant overlap between regions that gained H3K4me3 in *Setd1b* cKO oocytes and hypomethylated DMRs, whereas regions that lost H3K4me3 in the *Setd1b* cKO showed no apparent reciprocal gain in DNA methylation (Figure 4C, 4D, 4E). Therefore, these data again suggest that loss of SETD1B is accompanied by increased MLL2 action at certain sites, which show a reciprocal decrease in DNA methylation with gains of H3K4me3.

**Figure 4.**
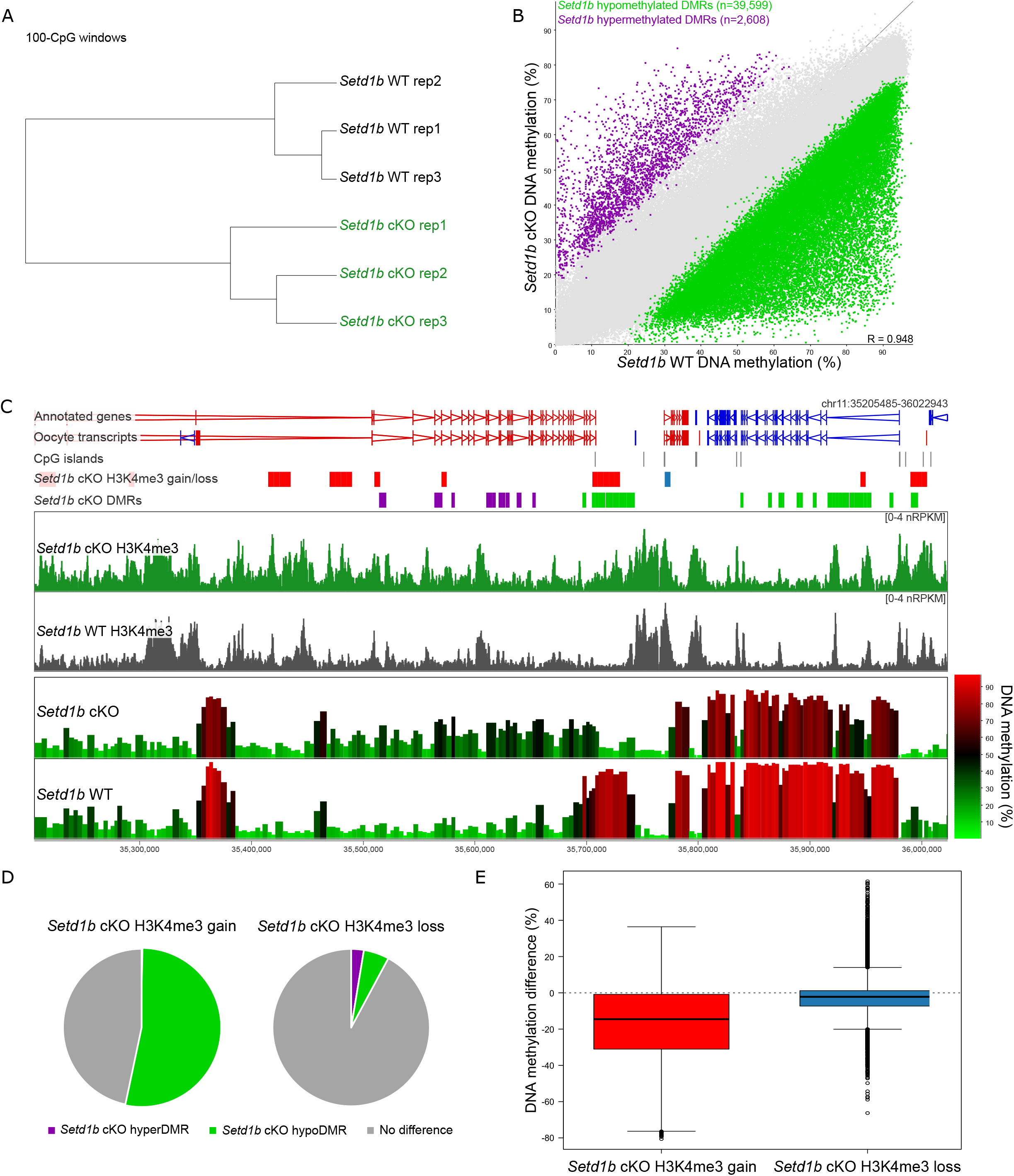
**(A)** Hierarchical cluster shows biological replicates for *Setd1b* cKO (N=3) and WT (N=3) GV oocyte DNA methylation patterns, quantitated as 100-CpG windows, with at least 10 informative CpGs (see Methods). **(B)** The scatterplot shows the DNA methylation in *Setd1b* cKO and WT GV oocytes across 100-CpG windows, with at least 10 informative CpGs in each replicate (N=424,105). Differentially methylated regions (DMRs) were identified using logistic regression (p<0.05 corrected for multiple comparisons) and a >20% difference in methylation. **(C)** The screenshot shows H3K4me3 enrichment of 1kb running windows with a 500bp step, and DNA methylation of 100-CpG windows with at least 10 informative CpGs per replicate, for *Setd1b* cKO and WT oocytes. Regions that gain or lose H3K4me3 in *Setd1b* cKO are shown in the labelled annotation tracks as red and blue bars, respectively. Regions that gain or lose DNA methylation (*Setd1b* DMRs) are shown in the labelled annotation tracks as purple and green bars, respectively. **(D)** The pie charts show the overlap between 5kb windows that gain (left) and lose (right) H3K4me3 in the *Setd1b* cKO with 100-CpG windows that are hypermethylated (HyperDMR) or hypomethylated (HypoDMR) in the *Setd1b* cKO (Chi-Square statistic, p<0.0001). **(E)** The boxplot shows the average difference in DNA methylation (*Setd1b* cKO – WT) of 100-CpG windows that overlap regions that gain and lose H3K4me3 in the *Setd1b* cKO (t-test, p<0.0001).

The mechanisms underlying the mutual exclusivity of H3K4me3 and DNA methylation are bidirectional, such that methylation of CpG dinucleotides can inhibit binding of the CxxC domain of MLL2 (7, 25, 26) and H3K4me3 impairs binding of the ADD domains of DNMT3A and DNMT3L (13, 14). In oogenesis, deposition of H3K4me3 and *de novo* methylation occur in parallel (40). Here, we sought to determine whether loss of SETD1B in oogenesis disrupts the balance between these two mechanisms. We reasoned that misdirected MLL2 H3K4 methylation activity in the *Setd1b* cKO would interfere preferentially with *de novo* DNA methylation occurring late in oogenesis. Using DNA methylation datasets from d12 and d15 GV oocytes (41), we evaluated the timing at which *Setd1b* hypomethylated DMRs normally acquire DNA methylation. Compared to a random set of methylated domains, *Setd1b* hypomethylated DMRs acquire methylation significantly later in oogenesis (Figure 5A, 5B). An alternative possibility is that delayed *de novo* DNA methylation in *Setd1b* cKO oocytes may permit the mistargeting of MLL2 to otherwise DNA methylated domains. We therefore compared the H3K4me3 enrichment between *Dnmt3a* and *Dnmt3b* conditional double knockout (termed *Dnmt3* cDKO) GV oocytes (40), which lack DNA methylation, and *Setd1b* cKO GV oocytes. H3K4me3 accumulation in *Dnmt3* cDKO oocytes is highly enriched among domains that would normally be DNA methylated (Figure 5C) (40), a trend that was partially recapitulated in the *Setd1b* cKO (Figure 5D, 5E). Together, these results indicate that SETD1B modulates the competition between MLL2 and *de novo* DNMTs for genomic targets in late oogenesis. In the absence of SETD1B, disruption of the balance between these mechanisms leads to impaired patterning of the oocyte epigenome.

**Figure 5.**
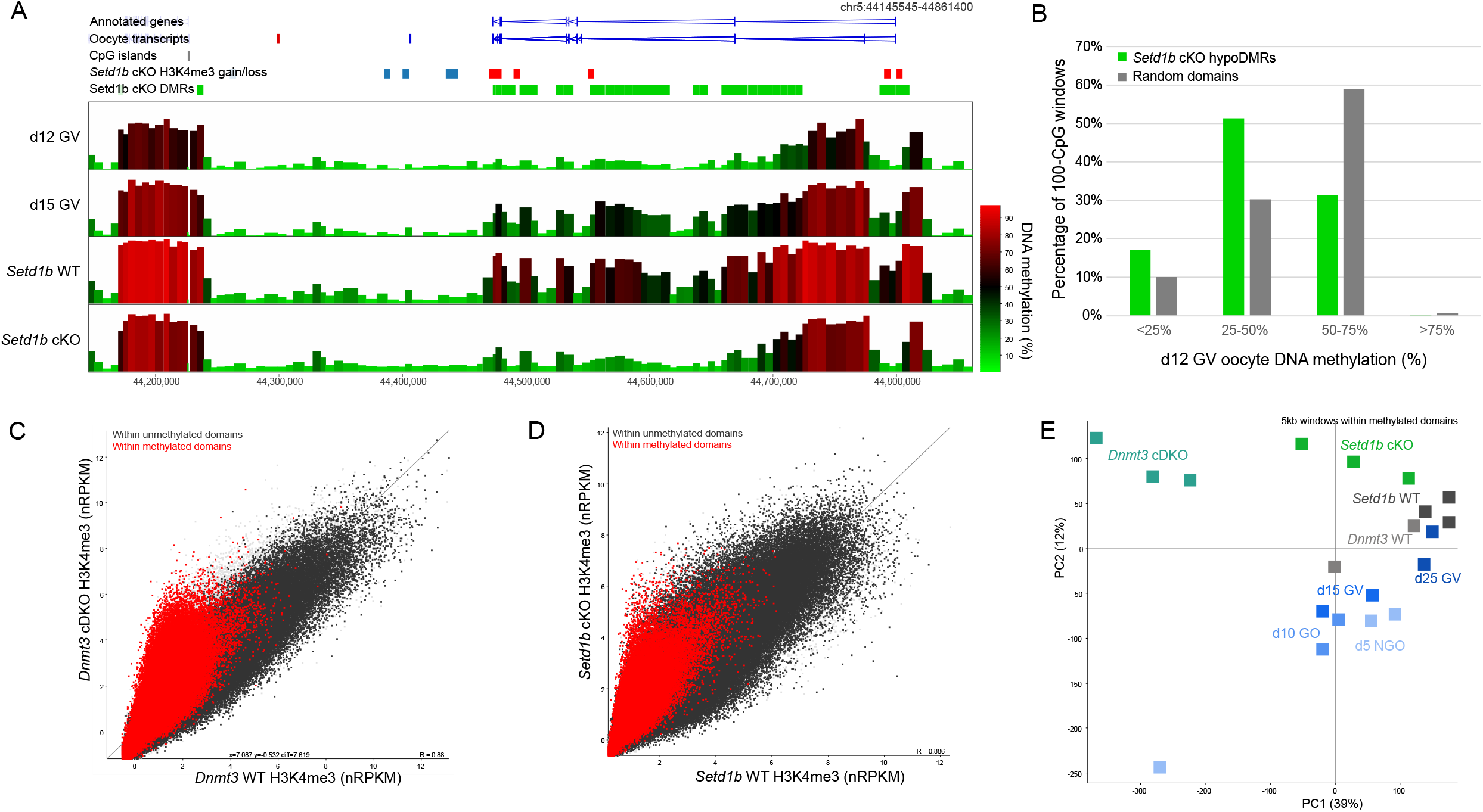
**(A)** The screenshot shows DNA methylation of 100-CpG windows for d12 GV, d15 GV, *Setd1b* WT GV (d21), and *Setd1b* cKO GV (d21) oocytes. Regions that gain or lose DNA methylation (*Setd1b* DMRs) are shown in the labelled annotation tracks as purple and green bars, respectively. Significant differentially enriched 5kb windows are shown in the *Setd1b* H3K4me3 gain/loss track, with red and blue bars showing windows that gain and lose H3K4me3 in *Setd1b* cKO, respectively. **(B)** The barplot shows the percentage methylation in d12 GV oocytes of *Setd1b* hypomethylated DMRs (hypoDMRs) and a random set of 100-CpG windows that fall within GV methylated domains (N=20,734 and 10,260, respectively). Chi-square analysis was used to compare the hypoDMRs to the random set (p<0.0001). **(C)** The scatterplot shows average normalised enrichment for H3K4me3 for 5kb running windows between *Dnmt3* cDKO (N=3) and WT (N=2) d25 GV oocytes. 5kb windows that fall within GV oocyte methylated domains are shown in red and unmethylated domains in dark grey. **(D)** The scatterplot shows average normalised enrichment for H3K4me3 for 5kb running windows between *Setd1b* cKO (N=3) and WT (N=3) d21 GV oocytes. 5kb windows that fall within GV oocyte methylated domains are shown in red and unmethylated domains in dark grey. (E) The principal component analysis plot shows H3K4me3 replicates for d5 NGO, d10 GO, d15 GV, d25 GV, *Setd1b* WT, *Setd1b* cKO, *Dnmt3* WT, and *Dnmt3* cDKO GV oocytes. The plot was generated using 5kb windows that were within GV oocyte methylated domains.

## Discussion

In this study, we provide the first detailed characterisation of epigenomic changes in *Setd1b* cKO oocytes. Previously, we discovered that MLL2 deposits broad, transcription-independent, H3K4me3 domains that are uniquely and abundantly found on the mouse oocyte genome (11, 40). However a subset of H3K4me3 peaks, associated with active promoters, were largely unaffected (40). Because SETD1B is essential for oogenesis (37), while SETD1A is not (27), we hypothesized that SETD1B would be responsible for H3K4me3 on these active promoters. Indeed, we find that a subset of active promoters lose both H3K4me3 and gene expression. Importantly, these genes are enriched for oocyte transcriptional regulators that are normally upregulated during oogenesis.

Unexpectedly, we observed a considerable reorganisation of genomic H3K4me3 involving both regional losses and gains. It appears that loss of SETD1B disturbs the genomic targeting of MLL2 and *de novo* DNMTs in late oogenesis, with increased H3K4me3 correlated with DNA hypomethylation across CpG-rich sequences. We speculate that this effect may be underpinned by either the increased stability or activity of MLL2 in *Setd1b* cKO oocytes, perhaps through increased availability of shared H3K4 methyltransferase cofactors upon the loss of SETD1B. These findings support a model whereby two complementary mechanisms target H3K4me3 to genomic loci, with SETD1B depositing H3K4me3 on active gene promoters supporting transcriptional activity, and MLL2 depositing H3K4me3 on unmethylated CpG-rich regions.

In addition to a strong association between H3K4me3 deposited by MLL2 and high CpG content in the oocyte, the association between MLL2 and CpG islands, including transcriptionally inactive promoters, has been observed in other contexts, such as embryonic stem cells (28, 52). Despite the relatively low affinity of CxxC domains for CpG dinucleotides (53, 54), MLL2 appears to uniformly occupy CpG islands (28) supporting the proposition that MLL2 has a general ability to associate with CpG islands in the naïve epigenome (36). Conversely, SETD1 complexes preferentially bind actively transcribed, H3K4me3-marked CpG island promoters (23, 55), in part due to the PHD finger of CXXC1 (23). Our data from oocytes supports the proposition that SETD1B is targeted to transcriptionally active promoters, likely through its CXXC1 subunit. Although MLL1 and presumably MLL2 contain a PHD finger that binds to H3K4me3 (56), the role of this interaction in their genomic distribution remains to be determined. Our observations in *Setd1b* cKO oocytes are consistent with our published findings that MLL2 activity in the oocyte is directed by binding to unmethylated CpGs rather than H3K4me3-marked regions (40).

As the predominant promoter-associated H3K4me3 methyltransferase in oocytes, loss of SETD1B was expected to provoke loss of gene expression. However, in concordance with observations in *Setd1b* cKO MII oocytes (37), we find the counterintuitive trend that more mRNAs are up-than down-regulated in the *Setd1b* cKO GV oocyte transcriptome. Furthermore, promoters of these upregulated genes show no gain in H3K4me3 in *Setd1b* cKO GV oocytes. This paradox may be explained by the observation that the downregulated genes are enriched for negative transcriptional regulators (37). In contrast to the upregulated genes, we found that downregulated genes are linked to reduced H3K4me3. Our data is consistent with a SETD1B/CXXC1-dependent deposition of H3K4me3 at these promoters early in oogenesis, suggesting that loss of H3K4me3 may precede a failure to induce expression of these genes in the *Setd1b* cKO oocytes. This impaired upregulation of gene expression in the absence of SETD1B, in turn, leads to substantial consequences for the transcriptional programme of the oocyte.

Consistent with the similar oocyte phenotypes observed between *Cxxc1* and *Setd1b* cKO (37, 38, 39), we also observe comparable molecular changes. These data suggest that CXXC1 and SETD1B are obligate partners in the deposition of H3K4me3 in oogenesis, and support the observation that, despite being present, SETD1A has a limited role (27).

In *Setd1b* cKO GV oocytes, almost as many sites gained H3K4me3, as lost H3K4me3. The gains were found on DNA hypomethylated, H3K27me3-marked, CpG-rich regions, indicating increased activity of MLL2. However, it is possible that a subset of site-specific changes at promoters and/or repetitive elements may be due to loss of transcriptional repressors in *Setd1b* cKO oocytes (37). Our observations are consistent with trends seen in *Cxxc1* cKO oocytes, which also showed a gain of H3K4me3 at H3K27me3-marked promoters (48). We speculate that in the absence of SETD1B there may be improved stability and/or abundance of MLL2-containing H3K4 methyltransferase complexes due to an increased availability of the core cofactors WDR5, RbBP5, ASH2L and DPY30. While this dynamic has not been widely discussed in the literature, it is consistent with trends observed in *Mll2* KO embryonic stem cells, where a reciprocal gain of H3K4me3 was observed at highly enriched promoters while lowly enriched bivalent promoters lost H3K4me3 (28). It would be difficult to test whether abundance of the core cofactors is rate limiting in oocytes, but could warrant future study in other cell contexts.

The redistribution of H3K4me3 in *Setd1b* cKO oocytes impacts the patterning of DNA methylation. The generalised loss of DNA methylation suggests a delay in the normal *de novo* methylation process in *Setd1b* cKO oocytes, although we did not detect reduced transcript abundance of the key *Dnmts*. In seeking to understand the altered DNA methylation landscape, we found that gains in H3K4me3 in *Setd1b* cKO oocytes partially recapitulate trends seen in oocytes lacking all DNA methylation. Furthermore, the high CpG content of regions that gain H3K4me3 in the *Setd1b* cKO may support a model where MLL2 is titrated away from CpG-poor inter-genic sites to more favourable CpG-rich sites when DNA methylation is delayed. Alternatively, because *de novo* methylation and nonpromoter H3K4me3 deposition are mutually exclusive and occur in parallel in growing oocytes (40), increased action or availability of MLL2 complexes could favour accumulation of H3K4me3 over domains that would otherwise be DNA methylated. Indeed, we observed that regions that acquire DNA methylation late in oogenesis were more likely to be impacted in the *Setd1b* cKO, suggesting that accumulating H3K4me3 may block late *de novo* DNA methylation. Together, our study demonstrates that MLL2 and *de novo* DNMTs compete for occupancy in late oogenesis and that loss of SETD1B disrupts the normal targeting of these enzymes.

We extend our previous proposition that MLL2 deposits H3K4me3 in broad domains to decoy promoter complexes and thereby contribute to global transcriptional silencing in late oogenesis (40). This enables the oocyte to be loaded with the factors and complexes involved in transcription ready for rapid activation upon removal of the broad domain H3K4me3 in the zygote. However, H3K4me3 at promoters does not need to be subject to removal. As identified here with SETD1B, the involvement of a second H3K4 methyltransferase focused on promoters allows the scope for these two H3K4me3 functions. Potentially, resistance to removal in the zygote suggests that the promoter H3K4me3 nucleosomes must be different from the broad domain H3K4me3 nucleosomes in some way.

In this study, we used ultra-low input sequencing methods to reveal molecular mechanisms underlying the targeting of H3K4me3 in oogenesis. Our findings help to further our understanding of how the H3K4 methyltransferases are functioning *in vivo* and co-ordinately contribute to the gene regulatory landscape in oocytes.

## Data availability

The sequencing data generated for this study has been deposited into the Gene Expression Omnibus database (GSE167987).

## Supplementary data

Supplementary Data are available at NAR Online.

## Acknowledgements

We would like to thank Heba Saadeh and Felix Krueger for their contributions in mapping and processing the post-bisulphite adaptor tagging data. We would like to thank the Sequencing and Bioinformatics facilities at the Babraham Institute. We thank the Biomedical Services of the Max Planck Institute of Molecular Cell Biology and Genetics, Dresden, for excellent service.

## Author contributions

A.F.S., A.K. and G.K. conceptualised the study. S.R., A.D., A.K., J.H. and C.H. collected the data. C.H., C.B., J.H., and S.A. performed data analysis. C.H., C.B., and S.A. generated manuscript figures. C.H. drafted the manuscript, with input from J.H., C.B., S.A., A.K., A.F.S. and G.K.

## Supplementary Item Legends

**Supplementary Table 1.** Raw and uniquely mapped read depths are provided for all sequencing libraries generated for this study. PBAT: post-bisulphite adaptor tagging

| Sample | Library | Raw Reads | Uniquely mapped |
| --- | --- | --- | --- |
| Setd1b WT rep1 | H3K4me3 ChIP-seq | 20,014,719 | 12,726,842 |
| Setd1b WT rep2 | H3K4me3 ChIP-seq | 40,103,864 | 10,579,220 |
| Setd1b WT rep4 | H3K4me3 ChIP-seq | 26,368,689 | 15,942,467 |
| Setd1b cKO rep1 | H3K4me3 ChIP-seq | 26,790,166 | 12,427,601 |
| Setd1b cKO rep3 | H3K4me3 ChIP-seq | 38,267,203 | 11,611,307 |
| Setd1b cKO rep4 | H3K4me3 ChIP-seq | 36,404,048 | 14,138,852 |
| Setd1b WT rep1 | PBAT | 65,496,468 | 29,075,293 |
| Setd1b WT rep2 | PBAT | 49,773,592 | 25,975,398 |
| Setd1b WT rep3 | PBAT | 60,201,429 | 32,757,844 |
| Setd1b cKO rep1 | PBAT | 69,046,033 | 29,903,629 |
| Setd1b cKO rep2 | PBAT | 63,538,005 | 33,224,984 |
| Setd1b cKO rep3 | PBAT | 68,637,891 | 34,873,524 |
| Setd1b WT rep1 | scRNA-seq | 16,915,059 | 9,329,696 |
| Setd1b WT rep2 | scRNA-seq | 17,712,949 | 9,726,867 |
| Setd1b WT rep3 | scRNA-seq | 14,765,556 | 8,006,481 |
| Setd1b cKO rep1 | scRNA-seq | 13,555,016 | 7,138,433 |
| Setd1b cKO rep2 | scRNA-seq | 16,236,758 | 8,997,586 |
| Setd1b cKO rep3 | scRNA-seq | 15,506,713 | 8,462,170 |
| Setd1b cKO rep4 | scRNA-seq | 11,803,207 | 6,063,243 |

**Supplementary Table 2.** Correlation matrix of R-values for all replicates of single cell RNA-seq. All oocyte transcripts were quantitated using log2RPM.

|  | Setd1b<br>cKO 1 | Setd1b<br>cKO 2 | Setd1b<br>cKO 3 | Setd1b<br>cKO 4 | Setd1b<br>WT 1 | Setd1b<br>WT 2 | Setd1b<br>WT 3 |
| --- | --- | --- | --- | --- | --- | --- | --- |
| **Setd1b**<br>**cKO 1** | 1.000 | 0.958 | 0.952 | 0.941 | 0.941 | 0.938 | 0.936 |
| **Setd1b**<br>**cKO 2** | 0.958 | 1.000 | 0.957 | 0.930 | 0.945 | 0.941 | 0.929 |
| **Setd1b**<br>**cKO 3** | 0.952 | 0.957 | 1.000 | 0.938 | 0.937 | 0.935 | 0.931 |
| **Setd1b**<br>**cKO 4** | 0.941 | 0.930 | 0.938 | 1.000 | 0.916 | 0.917 | 0.936 |
| **Setd1b**<br>**WT 1** | 0.941 | 0.945 | 0.937 | 0.916 | 1.000 | 0.960 | 0.948 |
| **Setd1b**<br>**WT 2** | 0.938 | 0.941 | 0.935 | 0.917 | 0.960 | 1.000 | 0.948 |
| **Setd1b**<br>**WT 3** | 0.936 | 0.929 | 0.931 | 0.936 | 0.948 | 0.948 | 1.000 |

**Supplementary Figure 1.**
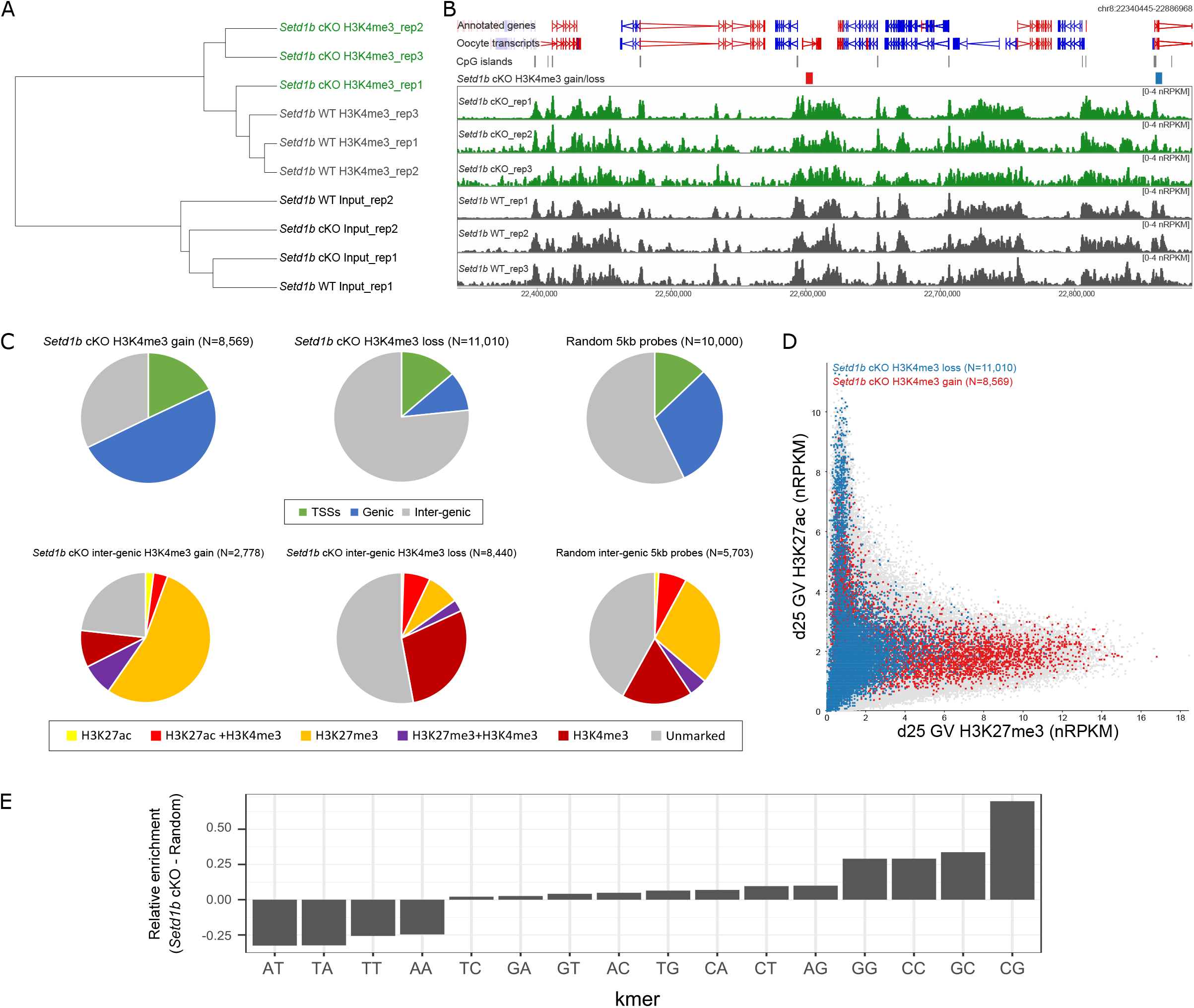
**(A)** Hierarchical clustering for H3K4me3 ChIP-seq replicates in *Setd1b* WT (N=3) and *Setd1b* cKO (N=3) GV oocytes, and 10% input replicates (N=4). Clustering was done using 5kb running windows and enrichment normalised RPKM. **(B)** The screenshot shows the normalised enrichment for H3K4me3 for 1kb running windows with a 500bp step in the *Setd1b* cKO and WT GV oocyte replicates. Significant differentially enriched 5kb windows are shown in the *Setd1b* gain/loss track, with red and blue bars showing windows that gain and lose H3K4me3 in *Setd1b* cKO, respectively. **(C)** Top: pie charts show the proportion of 5kb windows that gain (left) or lose (middle) H3K4me3 in the *Setd1b* cKO GV oocytes that overlap transcription start sites (TSSs) or genic regions, compared to a random set (right) of 5kb windows (p<0.0001 and p<0.0001, respectively, Chi-square). Bottom: pie charts show the proportion of the inter-genic 5kb windows that gain (left) or lose (middle) H3K4me3 in the *Setd1b* cKO GV oocytes that overlap combinatorial peaks for histone modifications in GV oocytes (Hanna et al., 2018), compared to the inter-genic random (right) 5kb windows (p<0.0001 and p<0.0001, respectively, Chi-square). **(D)** The scatterplot shows enrichment for H3K27ac and H3K27me3 for 5kb running windows (normalised RPKM) in d25 GV oocytes. The windows that gain and lose H3K4me3 in the *Setd1b* cKO are highlighted in red and blue, respectively. **(E)** The barplot shows the relative enrichment for dimer content in regions that gain H3K4me3 in the *Setd1b* cKO compared to a random set of 5kb probes.

**Supplementary Figure 2.**
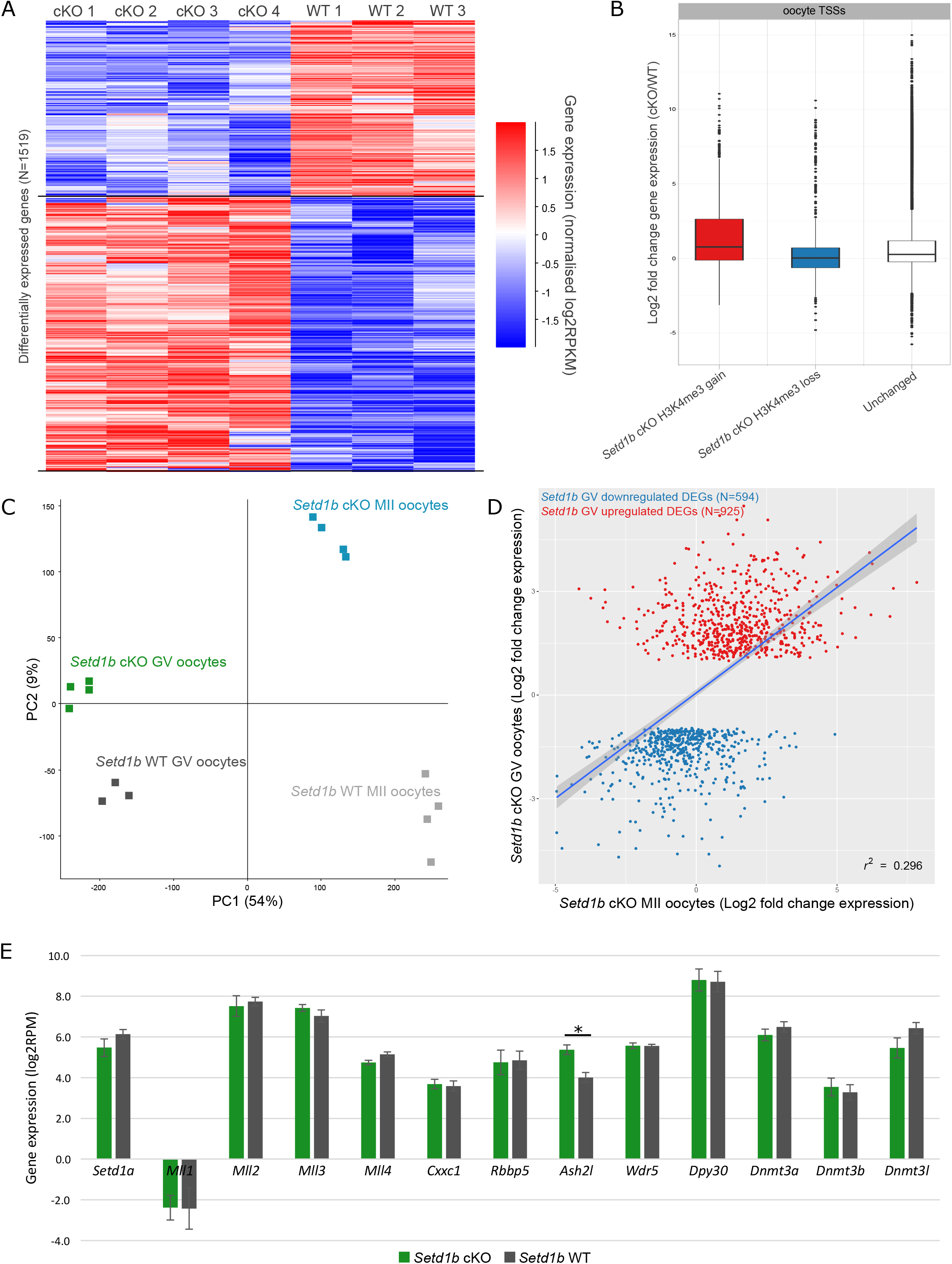
**(A)** The heatmap shows DEGs in the *Setd1b* cKO and WT GV oocyte replicates, quantitated as per gene-normalised log_2_RPKM. **(B)** The boxplot shows the log_2_ fold change in gene expression between the *Setd1b* cKO and WT for TSSs that overlap regions that gain (N=415), lose (N=914) or were unchanged (N= 26,428) for H3K4me3 in the *Setd1b* cKO GV oocytes. Pair-wise comparisons between gain and unchanged and loss and unchanged were done with two-tailed t-test (p=7.3E-8 and p=1.4E-14, respectively). **(C)** Principal component analysis shows scRNA-seq replicates for *Setd1b* cKO GV (N=4), *Setd1b* WT GV (N=3), *Setd1b* cKO MII (N=4), and *Setd1b* WT MII (N=4) oocytes. Expression of oocyte transcripts were quantitated using log_2_RPM. **(D)** The scatterplot shows log_2_ fold change (cKO/WT) in gene expression (RPKM) for *Setd1b* cKO DEGs in *Setd1b* cKO GV and MII oocytes. **(E)** The barplot shows the average gene expression for H3K4 methyltransferases (*Setd1a, Mll1, Mll2, Mll3, Mll4*), H3K4 methyltransferase complex subunits (*Cxxc1, Rbbp5, Ash2l, Wdr5, Dpy30*), and *de novo* DNA methyltransferases (*Dnmt3a, Dnmt3b, Dnmt3l*) in *Setd1b* cKO and WT GV oocytes. The whiskers show standard deviation between biological replicates, while the asterisks shows those genes that are significant DEGs.

**Supplementary Figure 3.**
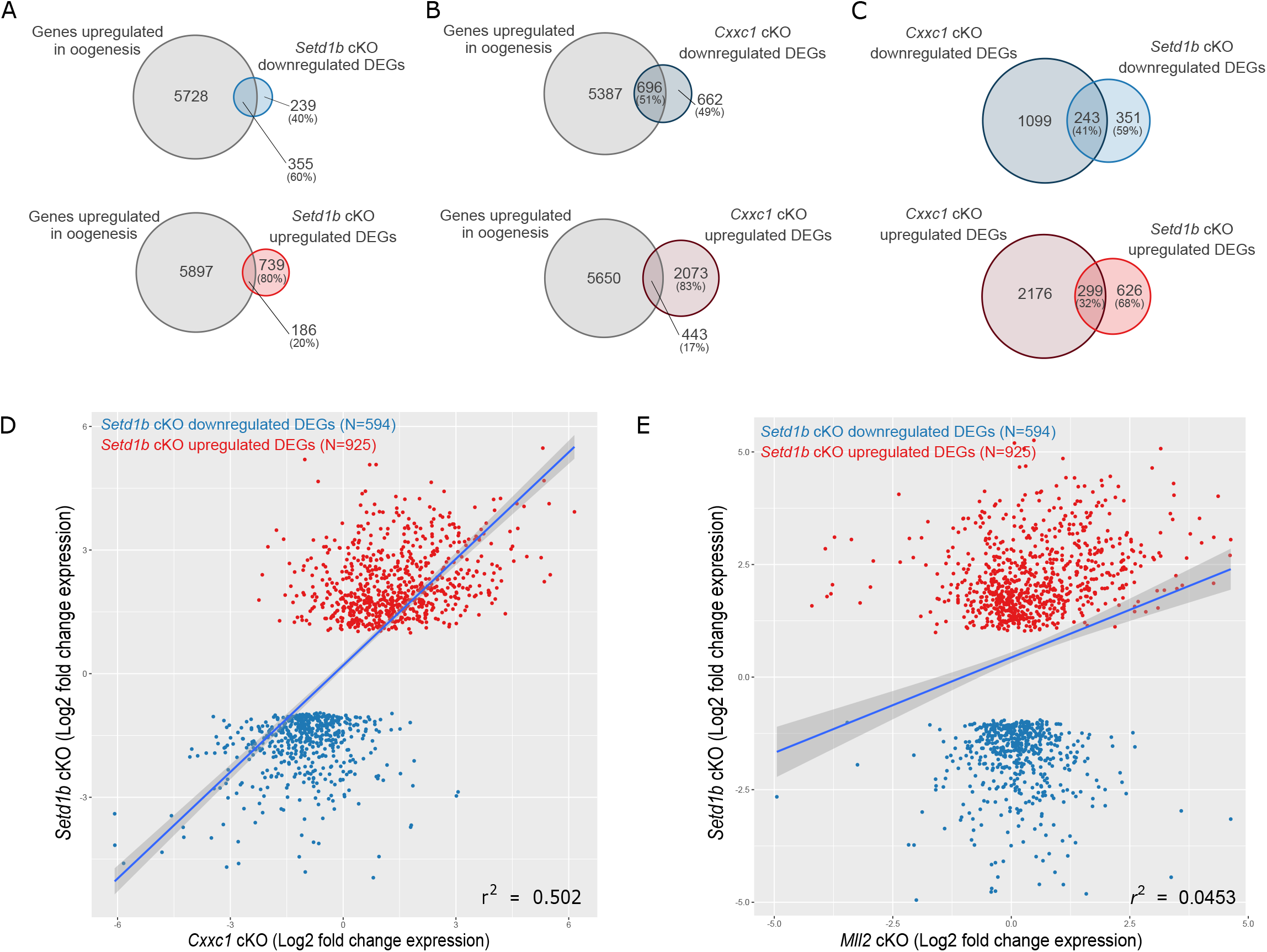
**(A)** The Venn diagrams show the overlap between genes upregulated in oogenesis and up- and downregulated DEGs in *Setd1b* cKO GV oocytes. **(B)** The Venn diagrams show the overlap between genes upregulated in oogenesis and up- and downregulated DEGs in *Cxxc1* cKO GV oocytes. **(C)** The Venn diagrams show the overlap between up- and downregulated DEGs in the *Cxxc1* cKO GV oocytes with up- and downregulated DEGs in *Setd1b* cKO GV oocytes. **(D)** The scatterplot shows the log_2_ fold change (cKO/WT) in gene expression (RPKM) for *Setd1b* cKO DEGs in the *Setd1b* cKO GV oocytes compared to the *Cxxc1* cKO GV oocytes. **(E)** The scatterplot shows the log_2_ fold change (cKO/WT) in gene expression (RPKM) for *Setd1b* cKO DEGs in the *Setd1b* cKO GV oocytes compared to the *Mll2* cKO GV oocytes.

**Supplementary Figure 4.**
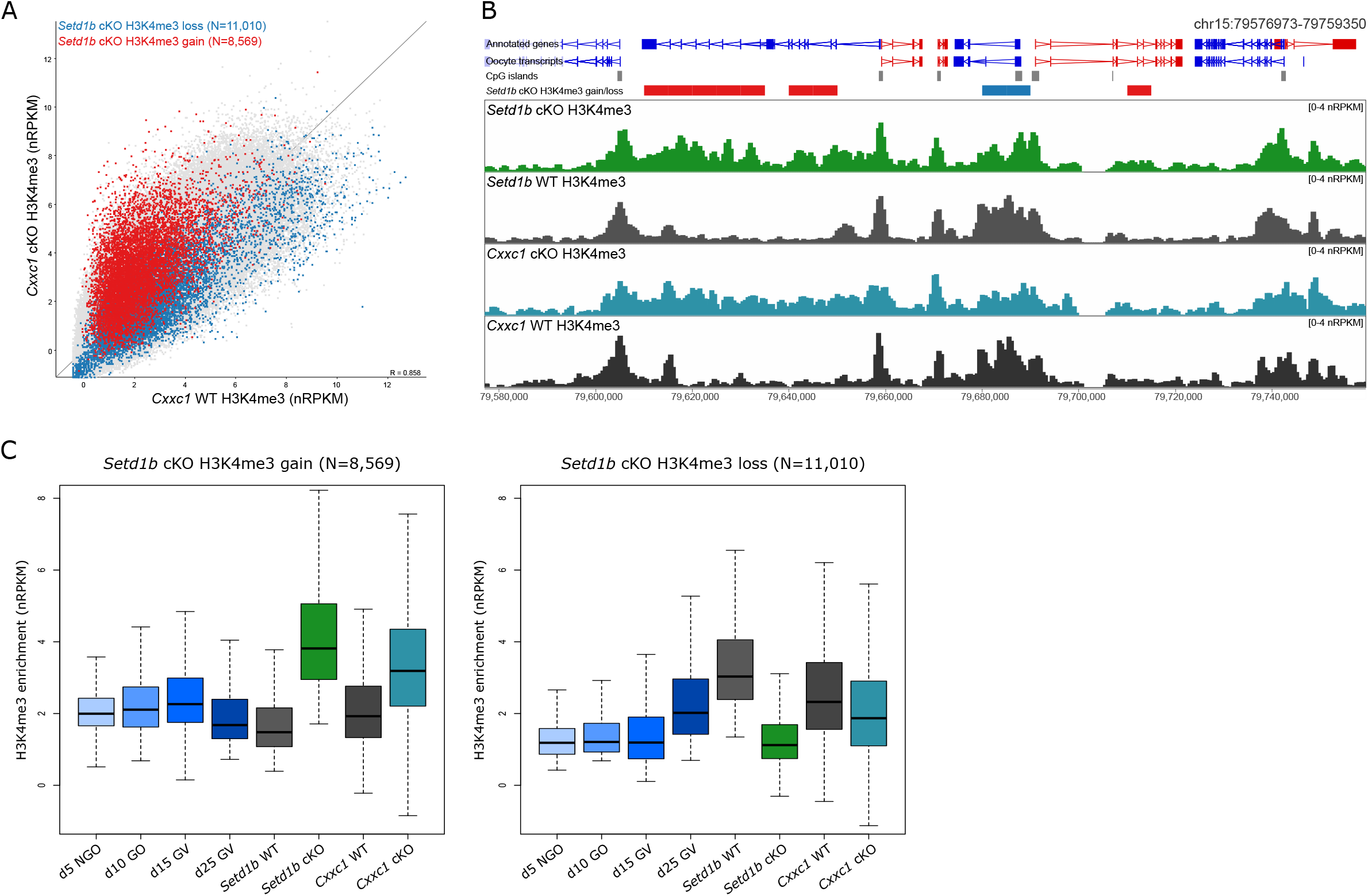
**(A)** The scatterplot shows average normalised enrichment for H3K4me3 for 5kb running windows (N=544,879) between *Cxxc1* cKO (N=2) and WT (N=2) GV oocytes. Differentially enriched windows identified in *Setd1b* cKO are shown in blue (loss) and red (gain). **(B)** The screenshot shows the normalised enrichment for H3K4me3 for 1kb running windows with a 500bp step in *Setd1b* cKO, *Setd1b* WT, *Cxxc1* cKO and *Cxxc1* WT GV oocytes. Significant differentially enriched 5kb windows are shown in the *Setd1b* gain/loss track, with red and blue bars showing windows that gain and lose H3K4me3 in *Setd1b* cKO, respectively. **(C)** The boxplot shows the H3K4me3 enrichment (normalised RPKM) in d5 non-growing oocytes (NGO), d10 growing oocyte (GO), d15 GV, d25 GV, *Setd1b* WT, *Setd1b* cKO, *Cxxc1* WT and *Cxxc1* cKO GV oocytes for 5kb windows that significantly gain (left) or lose (right) H3K4me3 in *Setd1b* cKO oocytes.

**Supplementary Figure 5.**
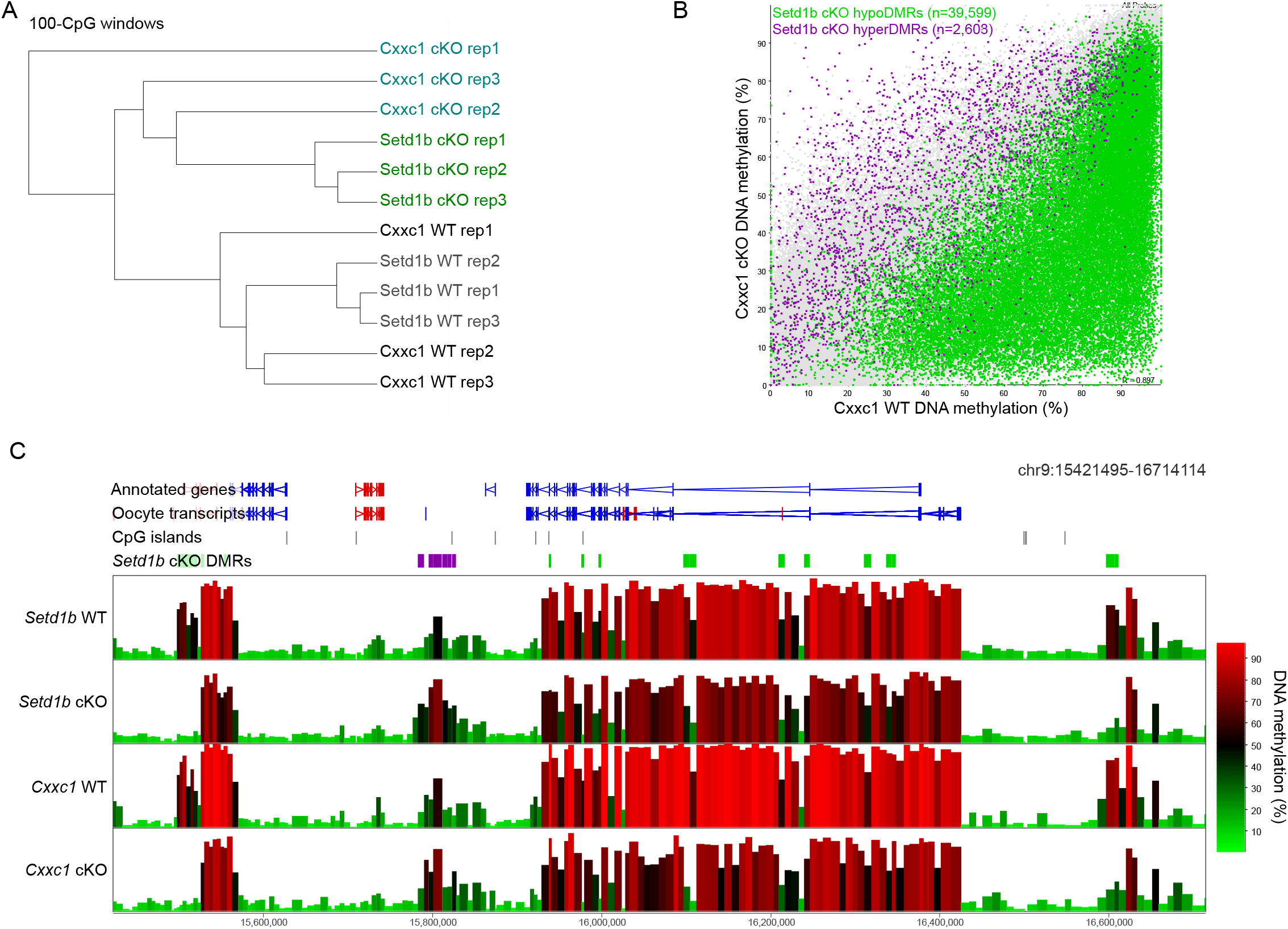
**(A)** Hierarchical clustering for replicates of DNA methylation in *Setd1b* WT, *Setd1b* cKO, *Cxxc1* WT and *Cxxc1* cKO GV oocytes. Clustering was done using 100-CpG windows (excluding the Y chromosome) with a minimum coverage of 10-CpGs. **(B)** The scatterplot shows the DNA methylation in *Cxxc1* cKO and WT GV oocytes across 100-CpG windows, with at least 10 informative CpGs in grouped replicates. *Setd1b* hypomethylated (green) and hypermethylated (purple) differentially methylated regions (DMRs) are highlighted. **(C)** The screenshot shows DNA methylation of 100-CpG windows with at least 10 informative CpGs per replicate for *Setd1b* cKO and WT oocytes and per grouped replicate for *Cxxc1* cKO and WT GV oocytes. Regions that gain or lose DNA methylation (*Setd1b* cKO DMRs) are shown in the labelled annotation tracks as purple and green bars, respectively.

